# Prior expectations guide multisensory integration during face-to-face communication

**DOI:** 10.1101/2025.02.19.638980

**Authors:** Giulia Mazzi, Ambra Ferrari, Maria Laura Mencaroni, Chiara Valzolgher, Mirko Tommasini, Francesco Pavani, Stefania Benetti

## Abstract

Face-to-face communication relies on the seamless integration of multisensory signals, including voice, gaze, and head movements, to convey meaning effectively. This poses a fundamental computational challenge: optimally binding signals sharing the same communicative intention (e.g. looking at the addressee while speaking) and segregating unrelated signals (e.g. looking away while coughing), all within the rapid turn-taking dynamics of conversation. Critically, the computational mechanisms underlying this extraordinary feat remain largely unknown. Here, we cast face-to-face communication as a Bayesian Causal Inference problem to formally test whether prior expectations arbitrate between the integration and segregation of vocal and bodily signals. Moreover, we asked whether there is a stronger prior tendency to integrate audiovisual signals that show the same communicative intention, thus carrying a crossmodal pragmatic correspondence. In a spatial localization task, participants watched audiovisual clips of a speaker where the audio (voice) and the video (bodily cues) were sampled either from congruent positions or at increasing spatial disparities. Crucially, we manipulated the pragmatic correspondence of the signals: in a communicative condition, the speaker addressed the participant with their head, gaze and speech; in a non-communicative condition, the speaker kept the head down and produced a meaningless vocalization. We measured audiovisual integration through the ventriloquist effect, which quantifies how much the perceived audio position is misplaced towards the video position. Bayesian Causal Inference outperformed competing models in explaining participants’ behaviour, demonstrating that prior expectations guide multisensory integration during face-to-face communication. Remarkably, participants showed a stronger prior tendency to integrate vocal and bodily information when signals conveyed congruent communicative intent, suggesting that pragmatic correspondences enhance multisensory integration. Collectively, our findings provide novel and compelling evidence that face-to-face communication is shaped by deeply ingrained expectations about how multisensory signals should be structured and interpreted.

**Author summary:** Face-to-face communication is complex: what we say is coupled with bodily signals, offset in time, which may or may not work in concert to convey meaning. Yet, the brain rapidly determines which multisensory signals belong together and which, instead, must be kept apart, suggesting that prior expectations play a crucial role in this decision-making process. Here, we directly tested this hypothesis using Bayesian computational modelling, which allows for isolating the contribution of prior expectations and sensory uncertainty on the final perceptual decision. We found that people have a stronger prior tendency to combine vocal and bodily signals when they convey the same communicative intent (i.e. the speaker addresses the observer concurrently with their head, gaze and speech) relative to when this correspondence is absent. Thus, the brain uses prior expectations to bind multisensory signals that carry converging communicative meaning. These findings provide key insight into the sophisticated mechanisms underpinning efficient multimodal communication.

## Introduction

Language evolved, is learned, and is often used in face-to-face interactions, where speech is coupled with multiple concurrent layers of bodily information, such as head movements and gaze shifts [1–5]. Some of these multisensory signals are directly relevant to the communicative exchange. For example, imagine you enter a busy restaurant and a waiter turns his body and head toward you, addresses you with a gaze shift and offers his help to find a seat. All these inputs, combined, signal the intention to communicate and thus carry congruent pragmatic meaning [1,4,6–9]. As such, it seems reasonable to assume that these signals belong to the same social act and should be integrated. Conversely, some multimodal signals may be tangential to the communicative exchange: they simply co-occur with each other but are not linked by the same underlying intention. For example, imagine that the waiter breaks into a brief cough: this signal is concurrent yet unrelated to the message being delivered and should therefore be segregated. Thus, temporal synchrony alone is not a reliable cue for integration because unrelated signals can be synchronous, while related signals can be temporally misaligned [4,10]. To further complicate the decision-making process, each conversational partner is taxed by fast turn-taking dynamics [11]. Yet, despite these critical constraints, we process multimodal communicative signals faster and more accurately than speech alone [12–17]. How do we achieve such an extraordinary feat so seamlessly and efficiently? Current behavioural evidence cannot address this question because it is descriptive: it indirectly suggests that multimodal signals facilitate human communication. Despite decades of research on multimodal integration and speech perception, the computational mechanisms underpinning seamless face-to-face communicative interactions remain largely unexplored.

Research on the normative principles of multisensory integration offers a powerful, computationally explicit platform for directly addressing this crucial, outstanding point. Importantly, we must first ask: what is the core computational challenge that we face when parsing the conversational scene? As exemplified above, face-to-face communication implies the fundamental binding problem of multisensory perception [5]: individuals must optimally integrate signals that come from a common source and segregate those from independent sources [18]. Critically, observers cannot directly perceive the underlying causal structure (i.e., the underlying causal relationship between signals). They must infer it based on the available crossmodal correspondences. Beyond the realm of face-to-face interactions, observers typically exploit a vast set of correspondences that includes temporal synchrony [19–26], spatial congruence [27–29], semantic correspondences [30–34] and synesthetic correspondences [35–37]. The (in)congruence of such cues modulates the strength of the observer’s common-cause assumption, that is the prior belief that signals should be integrated into one unified percept [37–42]. More generally, the common cause prior is supported by observers’ expectations [37,39], which in turn may depend on prior experience [43–47], training [48,49], ontogenetic [50] and phylogenetic [51] factors. Hierarchical Bayesian Causal Inference [18,52–56] provides a powerful normative strategy to arbitrate between the integration and segregation of multisensory inputs. For example, in a spatial localization task (e.g., judging the speaker’s position), observers should integrate auditory (e.g., voice position) and visual (e.g., body position) information according to the underlying causal structure, guided by the amount of audiovisual spatial disparity and thus the strength of the common cause prior [57–59]. Under a common source, observers should perform multisensory integration following the Forced Fusion principle, i.e. combining the unisensory components weighted by their relative sensory reliabilities [60–62]. Under independent sources, the unisensory signals should be processed independently. To compute a final response that accounts for causal uncertainty, observers should apply a final decision function, such as model averaging [52,53]: the estimates (e.g., spatial location) under the two potential causal structures are combined weighted by the posterior probabilities of these structures.

In face-to-face interactions, a crucial causal structure to infer is the speaker’s intention: for instance, whether the intention is to communicate (e.g. looking at the addressee while speaking) or otherwise (e.g. looking away while coughing). Audiovisual signals that are shared by the same communicative intention carry a potent crossmodal pragmatic correspondence [1,4,6–9]. Such correspondence may bolster the common cause prior and thereby guide the fast and efficient segmentation of the conversational scene into coherent, meaningful units. However, a simpler computational mechanism is also conceivable. Under the Forced Fusion account [60–62], observers mandatorily perform reliability-weighted integration without considering causal uncertainty. Here, information may be filtered according to its salience and relevance [63]: communicative bodily movements (e.g. facing and gazing at someone) are more perceptually salient [64–70] and more self-relevant [71,72] than non-communicative ones (e.g. looking away); thus, they may attract more attention and thereby guide the parsing of the conversational scene. Consequently, several key computational questions remain unanswered. First, during face-to-face communication, do people integrate multisensory signals following Bayesian Causal Inference or Forced Fusion principles? Second, is integration stronger for communicative than non-communicative signals? If so, why? On the one hand, head and gaze movements towards the observer may attract more attention than looking away, decrease the noise (i.e. increase the reliability) of the attended visual information and thereby increase its weight in the fusion process under both Forced Fusion and Bayesian Causal Inference accounts [73]. Alternatively, crossmodal pragmatic correspondences among communicative signals may reinforce the prior belief that these signals come from a common cause, as uniquely captured by Bayesian Causal Inference.

In two consecutive experiments, we formally tested these competing hypotheses by combining psychophysics and computational modelling of human behaviour. Our findings demonstrate that Bayesian Causal Inference best explains participants’ responses, demonstrating that prior expectations guide multisensory integration during face-to-face communication. Remarkably, multisensory integration was stronger for communicative than non-communicative signals due to an enhanced common cause prior, suggesting that pragmatic correspondences strengthen the a priori assumption that multisensory signals belong together. Collectively, the present findings offer novel and direct insight into the computational mechanisms driving the efficiency of multimodal communication.

## Experiment 1

### Materials and Methods

#### Participants

Participants were recruited using the recruitment platform and the social media pages of the University of Trento. The minimally required sample size was N=34, based on a-priori power analysis in G*Power [74] with a power of 0.8 to detect a medium effect size of Cohen’s d = 0.5 at alpha = 0.05. Accordingly, 34 participants were recruited and included in the final analyses (20 females; mean age 24, range 18-47 years). All participants were native speakers of Italian and reported normal or corrected-to-normal vision, normal hearing and no history of neurological or psychiatric conditions.

#### Ethics statement

The study was approved by the University of Trento Research Ethics Committee and was conducted following the principles outlined in the Declaration of Helsinki. All volunteers provided written informed consent before starting the experiment and received monetary reimbursement, course credits or a university gadget.

#### Stimuli

The stimuli consisted of audiovisual recordings of a female speaker uttering Italian words (for the communicative condition) or vocalizing the Italian vowel sound /a/ (for the non-communicative condition). The words were Italian common nouns divided into 22 groups according to their meaning. During the recording, the camera was placed in three different positions along the azimuth for the speaker to be on the right side, left side or at the centre of the screen, at respectively 9°, −9° and 0° of visual angle (Fig 1a). By recording from these three different positions, the head movements of the speaker were congruent with the position on the screen so that the participants would perceive the movements of the speaker as directed towards them. The participants were told that the speaker’s position would change to make the interaction task more challenging, similar to how communication can be more difficult when someone approaches us to initiate a communicative interaction in a noisy environment, like a busy restaurant. Words were recorded evenly between positions, and the vocalisation was recorded from all three positions. To address the participant in the communicative condition, each video started with the speaker looking down with her head bowed, then she lifted her head, looked at the camera and uttered the word [8]. Thus, each synchronous audiovisual stimulus exhibited a natural offset between the audio and video components, with bodily signals preceding the vocalization by approximately 500 milliseconds. Collectively, we introduced relevant linguistic content and bodily signals in the communicative condition to maximize its difference from the non-communicative condition. For all stimuli, the audio was recorded with a microphone facing the speaker, and background noise was removed afterwards using iMovie (macOS). The auditory stimuli were then convolved with spatially selective head-related transfer functions (HRTFs) and interpolated to obtain the desired locations using MATLAB (2021b) Audio Toolbox. The visual stimuli spanned 14 × 10 degrees visual angle and were presented on a black background. Guided by previous research [57,73], we opted for high visual reliability to ensure that observers could perform causal inference and successfully arbitrate between sensory integration and segregation. As a result, our study was optimised to assess causal inference on the observers’ reported sound perception.

**Fig 1.**
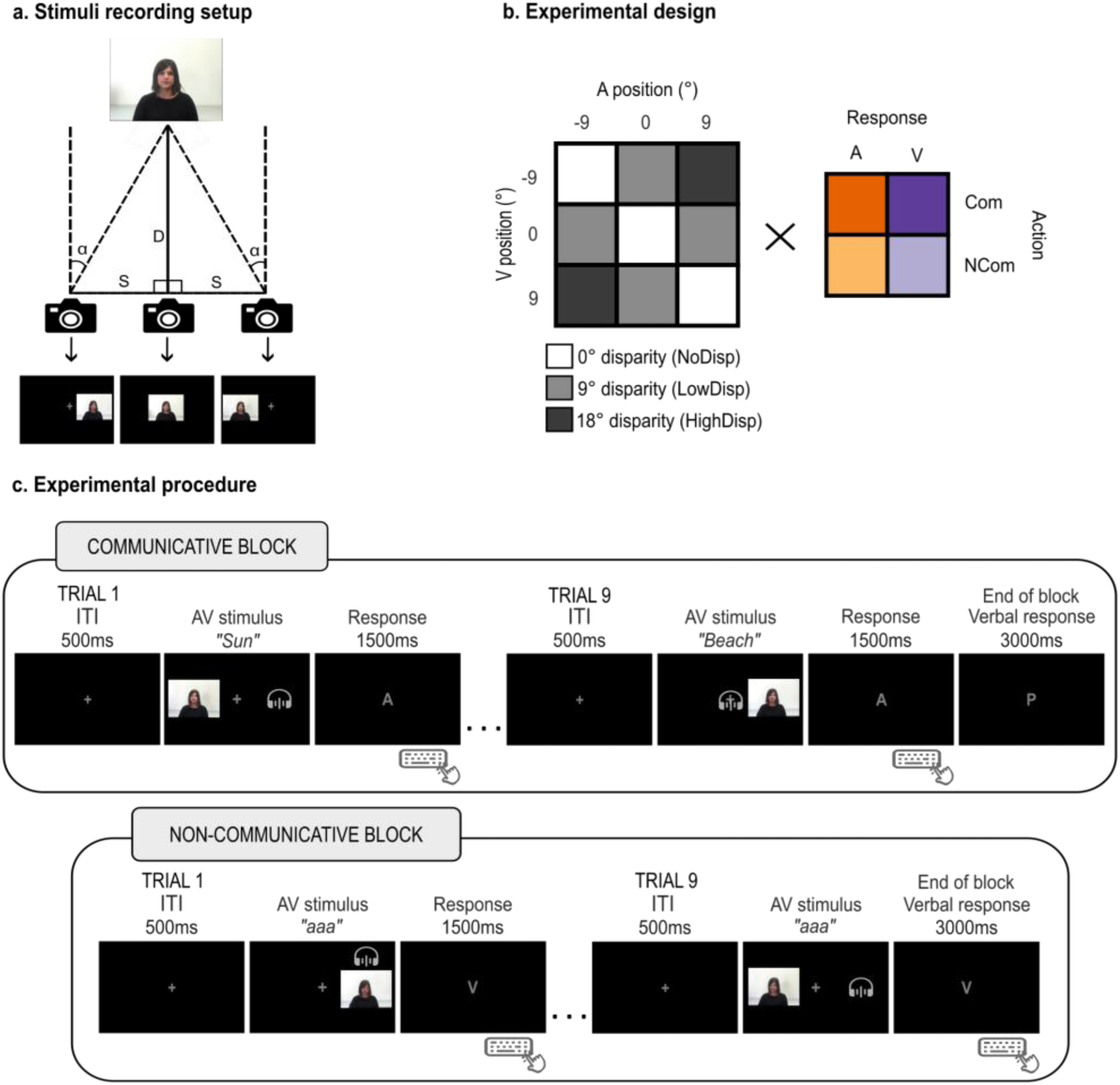
Stimuli, design and procedure. **a)** Visual stimuli were recorded from three different positions. D represents the distance between the speaker and the central camera (D = 150 cm). S represents the horizontal shift of the camera (23.6 cm) to record the speaker at an angle α (9 degrees visual angle), as defined by the formula: *S* = 2*D tan* (α/2). In the main experiment, videos with different viewing angles were presented on the side of the screen giving the impression that the speaker was facing the participant (e.g. videos recorded from the right side of the speaker were presented on the left side of the screen). **b)** We used a 3 (visual stimuli position) × 3 (auditory stimuli position) × 2 (action intention) × 2 (response modality) factorial design, resulting in 3 levels of spatial disparity (0, 9, 18 degrees of visual angle) and 36 total experimental conditions. **c)** The trial started with a 500 ms fixation cross at the centre of the screen, followed by the synchronous audiovisual stimulus. In communicative blocks, the stimuli were videos of the speaker turning towards the participant and uttering a word, which lasted on average 1500 ms (range 1000-2000 ms) depending on the length of the word. In non-communicative blocks, the stimuli lasted 500 ms in Experiment 1 and 1500 ms in Experiment 2. After the stimuli offset, a letter on the screen (‘A’ for auditory and ‘V’ for visual) cued participants to report the auditory or visual position using a specific keyboard key (1500 ms response window). At the end of communicative blocks, participants were prompted by the letter ‘P’ on the screen (for ‘parola’, i.e. ‘word’ in Italian) to say a word representing the general theme that connected the words in the block. The experiment consisted of 4 runs, each containing 18 blocks (9 communicative and 9 non-communicative) and 9 trials per block. The person depicted in the images is one of the authors.

#### Design and procedure

In a spatial ventriloquist paradigm, we presented synchronous audiovisual stimuli of a speaker uttering a word or producing a meaningless vocalisation. The auditory information (voice) and the visual bodily cues (video) were independently sampled from three positions along the azimuth (−9°, 0° or 9° of visual angles), generating 9 possible spatial combinations and 3 levels of spatial disparity (0°, 9°, 18° of visual angles; Fig 1b). Crucially, we manipulated the action intention: in half of the trials, the speaker directed her gaze toward the participant and uttered a word (communicative action); in the other half, a static frame of the speaker looking down was paired with a meaningless vocalisation (non-communicative action). After the presentation of each audiovisual stimulus, participants were instructed to report the position of either the visual or the auditory stimulus by a letter cue at the centre of the screen (‘A’ for audio and ‘V’ for video). The experiment was divided into blocks of 9 trials. The action intention (communicative vs. non-communicative) changed with each block, and the report modality (auditory vs. visual) changed every two blocks. Within each block, each spatial combination was presented once in a randomized order. In summary, the experiment conformed to a 3 (visual stimuli positions: −9°; 0°; 9° of visual angles) × 3 (auditory stimuli position: −9°; 0°; 9° of visual angles) × 2 (action intention: communicative; non-communicative) × 2 (response modality: auditory; visual) factorial design, for a total of 36 experimental conditions. Each participant completed 4 runs, with 9 trials/block and 18 blocks/run, thus generating 18 trials/condition and 648 trials in total. Each trial started with a 500 ms grey fixation cross in the middle of the screen on a black background, followed by synchronous auditory and visual stimuli. The fixation cross was present for the entire duration of the stimuli and participants were instructed to fixate it throughout the experiment. The communicative stimuli lasted on average 1500 ms (range 1000-2000 ms) considering the bodily movements preceding the utterance and the duration of the word. The non - communicative stimuli had a fixed duration of 500 ms. This shorter duration was chosen to optimize the length of the experiment for subsequent neuroimaging studies. After the stimuli disappeared, the letter ‘A’ or ‘V’ replaced the fixation cross on the screen to indicate the required response modality. This letter remained on the screen for 1500 ms, during which participants had to respond by identifying the location of the target stimuli using a specific keyboard key (left, down and right arrows for left, central and right locations respectively).

To increase ecological validity and ensure that participants focused not only on the spatial position of the stimuli but also on the communicative aspect and the semantic meaning of the words – similar to a real-life face-to-face conversation – they were informed that the person in the videos was interacting with them via a real-time online connection. Moreover, all nine words within each communicative block were connected by a common theme that the speaker aimed to convey to the participant (for example, “sun”, “holidays” and “beach” were linked by the theme “summer”). At the end of each communicative block, participants were prompted by a letter cue in the centre of the screen to guess the theme.

#### Experimental setup

The experiment was presented with Psychtoolbox (version 3.0.18) running in MATLAB R2021a on a Windows machine (Microsoft 10). Auditory stimuli were presented via headphones (Sannheiser HD580 precision) and visual stimuli were presented on a monitor (1920×1089 pixels resolution; 100 Hz frame rate). Participants sat in a dimly lit cubicle in front of the computer monitor at a viewing distance of 80 cm with their heads positioned on a chin rest to minimise movements. To ensure participants maintained central fixation, we monitored and recorded their ocular movements using an eye tracker device (EyeLink® 1000, version 4.594, SR Research Ltd.). At the end of each trial, participants responded using the keyboard’s left, down or right arrow keys.

#### Statistical analyses - Overview

We limited our analyses to trials without missed responses (i.e. no answer within a 1.5-second response time window), premature responses (RT < 100 ms) or response outliers (| RT | > 3 SD from the across-condition median RT of each participant). Only a few trials were discarded (5.7% ± 0.6% [across participants mean ± SEM]). Further, we excluded trials without central fixation during stimuli presentation. Significant saccades were defined as eye movements with an amplitude greater than 7° (half the width of the video stimuli, hence exceeding the video boundaries). Participants successfully maintained central fixation with only a few rejected trials (3.78% ± 0.42% [across participants mean ± SEM]).

In the following, we describe the main analyses: audiovisual weight index (*w_AV_*) and Bayesian modelling. Although not central to the present study, we additionally examined response times (see Supplementary materials).

#### Audiovisual weight index w_AV_

To evaluate the effect of auditory and visual signal locations on observers’ reported auditory (or visual) locations, we calculated an audiovisual weight index (*w_AV_*) for the trials where the auditory and visual signals were spatially incongruent (i.e., AV spatial disparity greater than 0). The *w_AV_* index is defined as the difference between the reported location in an incongruent trial (e.g. when the visual stimulus is at 0° and the auditory stimulus at 9°) and the average reported location in audiovisual congruent trials (e.g. when audiovisual stimuli are at 9°), scaled by the difference between the average reported locations in the two respective congruent condition (e.g. when audiovisual stimuli are at 0°, and when audiovisual stimuli are at 9°) [73]:

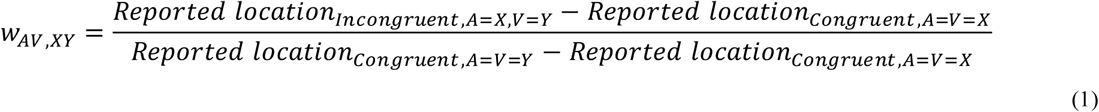

The denominator acts as a scaling operator for all conditions at a particular level of spatial disparity; to increase its estimation precision, we computed it using the average reported location in audiovisual congruent trials pooled over all conditions across all participants (A=V=-9°: −8.589°; A=V=0°: −0.667; A=V=9°: 8.208°). Notice that using the perceived locations of the congruent conditions instead of the true stimuli locations accommodates participants’ response biases. Under the assumption of comparable biases across all conditions, the *w_AV_* index represents the relative influence of the auditory and visual positions on the reported position: a value of 0 indicates that the observer’s report relies completely on the auditory signal position; a value of 1 indicates that the observer’s report relies completely on the visual signal position.

We computed the *wAV* index for each participant and experimental trial using Eq. 1 and we then averaged the values across all AV combinations at a certain level of spatial disparity (9°: low; 18°: high). Hence, we analysed the *wAV* index in a 2 (response modality: auditory; visual) × 2 (spatial disparity: low; high) × 2 (action intention: communicative; non-communicative) factorial design. To avoid making parametric assumptions, we evaluated the main effects of response modality, spatial disparity, action intention and their interactions using two-tailed permutation testing at the between-subject level (4,096 cases). For effect sizes, we report the difference between the across-participants’ mean empirical effect and the mean of the nonparametric null distribution and 95% Cis [73].

#### Bayesian modelling

Combining psychophysics and Bayesian modelling, we characterized the computational principles by which the observer integrates multisensory signals during face-to-face communication. In the following, we first describe the Bayesian Causal Inference model from which we derive the Forced Fusion model as a special case (see [52] for further details). Second, we describe how action intention may affect multisensory processing.

The Bayesian Causal Inference (BCI) model formalizes how an ideal observer should combine information deriving from different sensory modalities while considering the uncertainty about the underlying causal structure, i.e. whether sensory events are generated by a common cause or separate causes (Fig 2a).

**Fig 2.**
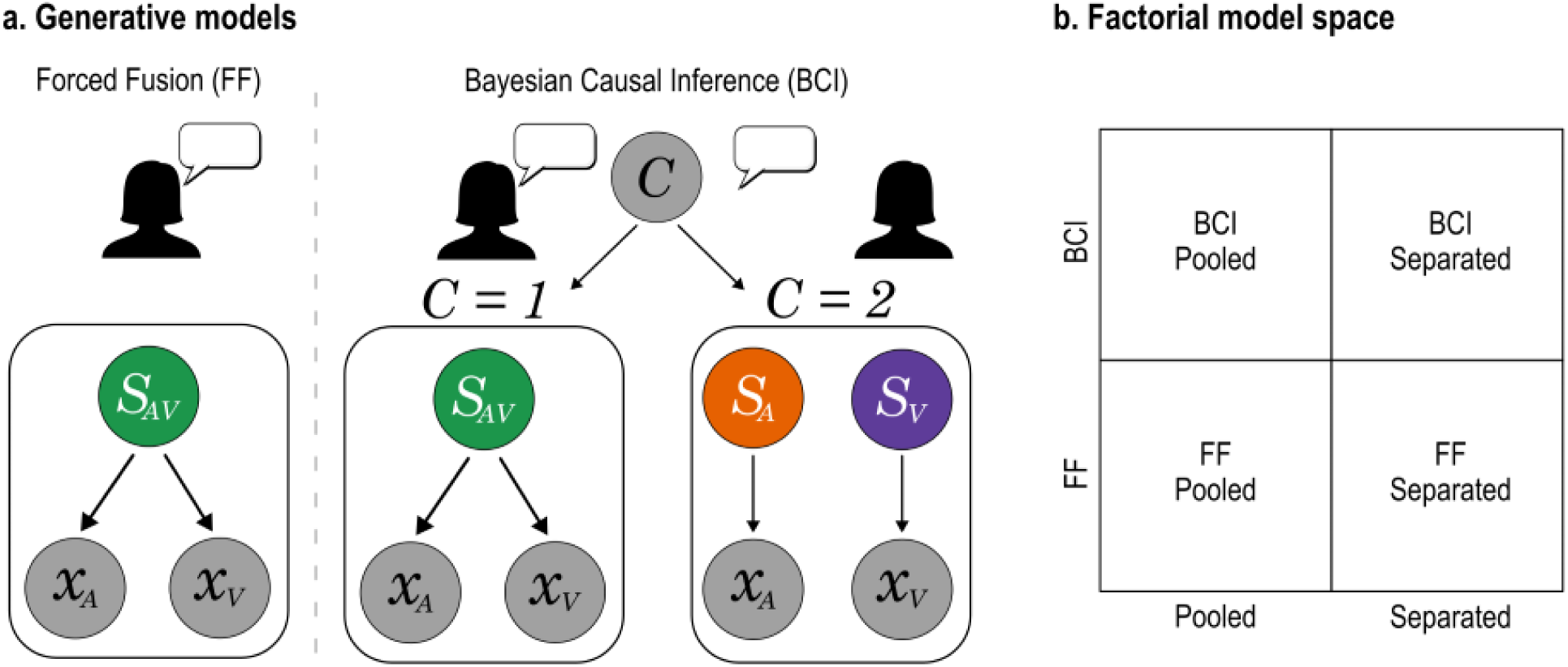
Bayesian Modelling. **a)** Generative models of Forced Fusion (FF) and Bayesian Causal Inference (BCI). For FF, a single sourcemandatorilygeneratesauditoryandvisualsignals. Instead, BCI explicitly models the two causal structures, i.e., whether auditory and visual signals come from one common cause (*C=1*) or separate causes (*C=2*). The FF model assumes that the auditory (*x*_*A*_) and the visual (*x*_*V*_) signals originate from the same source (*S*_*AV*_). The BCI model accounts for causal uncertainty, i.e. the possibility that the auditory (*x*_*A*_) and the visual (*x*_*V*_) signals could originate from one common (*C=1*) or two independent (*C=2*) causes. Due to its model architecture (see section “Bayesian modelling”), only BCI models accommodate spatial disparity and response modality effects. **b)** To determine whether a BCI or FF model best explained each participant’s localisation responses, and to evaluate the modulatory influence of action intention (communicative vs. non-communicative), we performed Bayesian model comparison in a 2 (BCI vs FF) × 2 (Pooled vs Separated action intention conditions) factorial model space.

Briefly, the BCI model assumes that the common (*C=1*) and independent (*C=2*) causes are sampled from a binomial distribution defined by the common cause prior *p*_common_, which represents the prior probability that the multisensory signals belong to the same source [52]. In other words, *p*_common_ quantifies the observer’s unity assumption, that is the prior belief that the signals should be integrated into one unified percept [37–42]. Such common cause prior is likely supported by observers’ expectations [37,39], which in turn may depend on prior experience [43–46], training [48,49], ontogenetic [50] and phylogenetic [51] factors. In the context of spatial localization, we know that auditory and visual stimuli that are spatially congruent tend to originate from the same source. Accordingly, audiovisual spatial disparity impacts the observer’s unity assumption: integration increasingly breaks down at higher disparity levels, where auditory and visual signals are less likely to originate from a common source. The computational architecture of the BCI model directly accounts for this effect [52]. Under the assumption of a common cause, the position of the source *S* is drawn from the prior distribution *N*(0, σ_*P*_), thereby assuming a central bias [57,75–77] with a strength represented by σ_*P*_. Under the assumption of separate causes, the positions *S*_*V*_ and *S*_*A*_ are drawn independently from the same distribution *N*(0, σ_*P*_). We then assume that auditory and visual signals are corrupted by independent Gaussian noise of standard deviations σ_*V*_ and σ_*A*_; thus, the events *x*_*V*_ and *x*_*A*_ are drawn respectively from the independent Gaussian distributions *N*(*S*_*V*_, σ_*V*_) and *N*(*S*_*A*_, σ_*A*_) [52]. Accordingly, the generative model employs four free parameters: common cause prior probability (*p*_common_), standard deviation of the spatial prior (σ_*P*_), and standard deviation of the auditory and visual representations (σ_*A*_ and σ_*V*_).

The posterior probability of the underlying causal structure is inferred by combining the common cause prior *p*common with the sensory evidence according to Bayes’ rule:

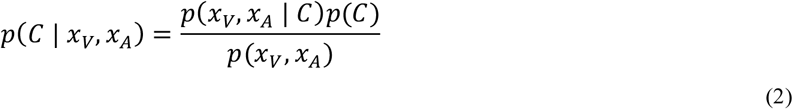

Depending on the causal structure, the brain generates the respective perceptual estimates by linearly weighting the signals *x*_*V*_ and *x*_*A*_ by the reliability of the respective sensory modality, i.e. the inverse of their variances. In the case of two independent sources, the optimal visual and auditory estimates are:

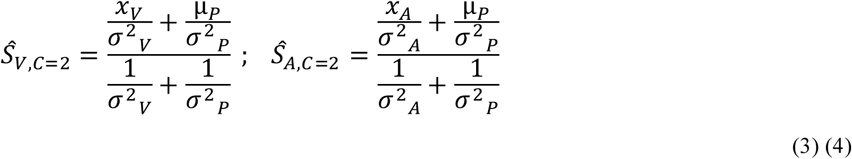

In the case of a common cause, thus *S*^_*V*_ = *S*^_*A*_, the optimal estimate is:

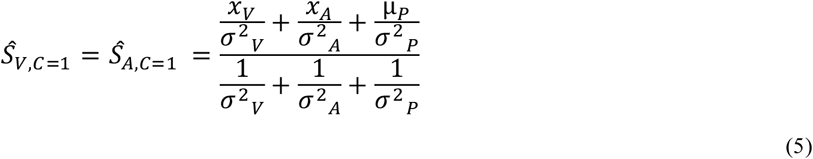

To compute the final estimate of the auditory and visual locations under causal uncertainty, the brain may use various decision functions, such as “model averaging”, “model selection” and “probability matching” [78]. In line with previous work [58,59,73,79], we focused on model averaging, which integrates the estimates of each causal structure weighted by the posterior probability of the respective structure. Accordingly, the final perceptual estimates of the auditory and visual source locations are computed as:

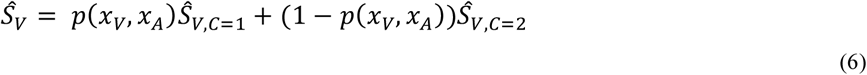

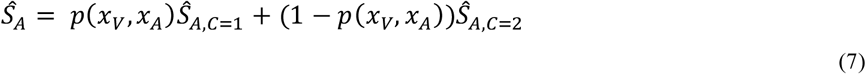

Thus, the Bayesian Causal Inference model relies on three spatial estimates, Ŝ_*AV,C*=1_, Ŝ_*V,C*=2_ and Ŝ_*A,C*=2_, which are combined into one final estimate according to task relevance (Ŝ_*A*_ for auditory response, Ŝ_*V*_ for visual response). In other words, the final perceptual estimate reflects the flexible readout of internal estimates according to current task demands [73]. Collectively, two effects naturally arise from the BCI model architecture: first, integration weakens when auditory and visual signals are sampled from more distant locations (effect of spatial disparity); second, task relevance dictates which internal perceptual estimates are combined to generate the final perceptual response (effect of response modality).

The Forced Fusion (FF) model can be considered a particular scenario of the Bayesian Causal Inference are therefore mandatorily integrated (i.e., p_common_ = 1). The final spatial audiovisual estimate Ŝ_*AV*_ is directly computed as a reliability-weighted linear average of the two unisensory estimates as described by Eq. 5 [80]. Thus, the FF model includes only three free parameters: standard deviation of the spatial prior (σ_*P*_), and standard deviation of the auditory and visual representations (σ_*A*_ and σ_*V*_). As a result, the FF model architecture does not accommodate the effects of spatial disparity or response modality.

To determine whether a BCI or FF model would best explain each participant’s localisation responses, and to evaluate the modulatory influence of action intention (communicative vs. non-communicative), we performed Bayesian model comparison in a 2 (BCI vs FF architecture) × 2 (Pooled vs Separated action intention conditions) factorial model space (Fig 2b). Thus, we compared 4 models: (i) a BCI model that does not account for action intention, (ii) a BCI model with separate parameters for each action intention condition, (iii) an FF model that does not account for action intention, and (iv) an FF model with separate parameters for each action intention condition. By comparing FF vs BCI models, we evaluated whether observers always integrate auditory and visual stimuli regardless of causal uncertainty (in line with FF), or if they combine the two alternative perceptual estimates weighted by the posterior probabilities of the respective causal structure (in line with BCI) as described by Eq. 6 and 7. This would confirm that prior expectations arbitrate between the integration and segregation of vocal and bodily signals during face-to-face communication [4,5]. By comparing Pooled vs Separated models, we evaluated whether observers showed a stronger tendency to integrate vocal and bodily information for communicative than non-communicative signals. Crucially, by comparing the BCI versus FF Separated models we characterized the exact computational mechanism by which communicative signals may increase multisensory integration. On the one hand, the perceptual characteristics of the audiovisual stimuli may influence the reliabilities of the sensory information before integration in both the FF and the BCI models. Specifically, the speaker may capture the observer’s attention more effectively by addressing them with head and gaze (communicative condition) as opposed to keeping the head down (non-communicative condition), thereby decreasing the noise (i.e. increase the reliability) of the attended visual information [73]. This would be captured by a decrease of σ_*V*_ in the communicative than non-communicative condition, which would increase the visual weight during the fusion process under the communicative condition. Alternatively, the shared communicative nature of the video and audio stimuli may reinforce the prior belief that the two signals come from a common cause, as captured by an increase of p_common_ in the BCI model for communicative than non-communicative trials. The factorial model comparison allowed us to directly arbitrate between these competing hypotheses.

To summarize, the BCI Pooled model employs four free parameters (*p*_*common*_, σ_*P*_, σ_*V*_ and σ_*A*_), which are doubled in the BCI Separated model (communicative condition: *p*_*common Com*_, σ_*P Com*_, σ_*V Com*_ and σ_*A Com*_; non-communicative condition: *p*_*common NCom*_, σ_*P NCom*_, σ_*V NCom*_ and σ_*A NCom*_). The FF Pooled model includes three free parameters (σ_*P*_, σ_*V*_ and σ_*A*_). Following the same logic as above, these three parameters are doubled in the FF Separated model that considers the two action intentions separately (communicative condition: σ_*P Com*_, σ_*V Com*_ and σ_*A Com*_; non-communicative condition: σ_*P NCom*_, σ_*V NCom*_ and σ_*A NCom*_).

We fitted each model to the individual participants’ localisation responses based on the predicted distributions of the spatial estimates. The distributions were obtained by marginalising over the internal variables not accessible to the experimenter (*x*_*V*_ and *x*_*A*_), which were simulated 10,000 times for each of the 36 conditions (see ‘Experimental Design’), and inferring the spatial estimates from Eqs. 2–7 for each simulated *x*_*V*_ and *x*_*A*_. Thus, we obtained a histogram of the auditory and visual responses predicted by each model for each condition and participant; based on these, we applied the multinomial distribution to compute the probability of each participant’s counts of localisation responses [73]. This gave us the likelihood of each model given the participants’ responses. Assuming independence of conditions, we summed the log-likelihoods across conditions. To obtain maximum likelihood estimates for the parameters of the models, we used a nonlinear simplex optimisation algorithm as implemented in MATLAB’s *fminsearch* function (MATLAB R2022b), after initialising the parameters in a prior grid search.

We firstly assessed the model fit for the data using the coefficient of determination R^2^, defined as:

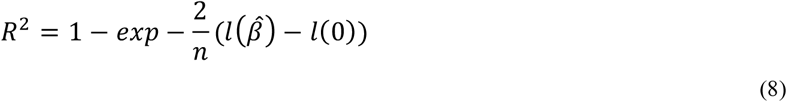

where *l*(0) and *l*(β^^^) represent respectively the log likelihood of the null model and the fitted model, and *n* is the number of data points. As the null model, we assumed a random response between the three possibilities (i.e. left, centre, right), hence a discrete distribution with a probability of 0.33. As in our case the models’ responses were discretized to relate them to the three discrete response options, and the coefficient of determination was scaled (i.e., divided) by the maximum coefficient defined as:

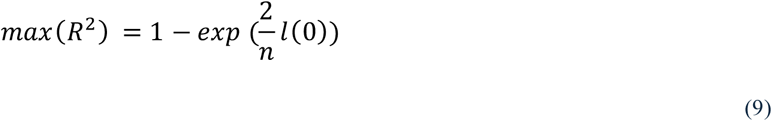

To identify the model that fits the localisation responses best, we compared the models using the Bayesian Information Criterion as an approximation to the log model evidence [81]:

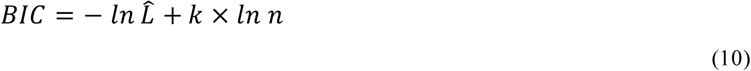

where *L^^^* represents the likelihood, *k* the number of parameters and *n* the number of data points. We then used SPM12 software (Statistical Parametric Mapping, Functional Imaging Laboratory, UCL) to perform random - effect Bayesian model selection at the group level [82]. This allowed us to estimate the protected exceedance probability that one model explains the data better than any other models and above the chance level, as quantified by the Bayesian Omnibus Risk (BOR). Lastly, for the winning model (i.e. the BCI Separated model), we evaluated pairwise parameter changes as a function of action intention (communicative vs. non - communicative). To refrain from parametric assumptions, we employed non-parametric pairwise tests (Wilcoxon signed-rank tests).

## Results

### Audiovisual weight index w_AV_

To evaluate *whether* response modality, spatial disparity and action intention influenced multisensory integration, we examined their main effects and interactions on the audiovisual weight index *w_AV_*, which quantifies the impact of the visual and auditory signals’ positions on participants’ spatial reports. Results are shown in Fig 3 (top panels) and summarized in Table 1 (see also S1 Table for comprehensive descriptive statistics). We found a significant main effect of response modality (*p* < 0.001, es = 0.83, CI(95%) = [0.79, 0.86]), indicating that participants relied more on the auditory signal when reporting the auditory position, and relied more on the visual signal when reporting the visual position (Fig 3a). Thus, task relevance influenced participants’ localization responses, corroborating the presence of a flexible readout mechanism of internal perceptual estimates according to current task demands [58,73]. Additionally, we found a significant response modality × spatial disparity interaction (*p* = 0.000, es = 0.09, CI(95%) = [0.08, 0.11]). Post hoc permutation testing showed that the *w_AV_* index depended on spatial disparity when participants reported the auditory position (*p* = 0.000, es = −0.09, CI(95%) = [−0.11, −0.07]), reflecting a greater influence of the visual stimuli on spatial localization when audiovisual signals were closer in space. These results confirm that integrating signals from different sensory modalities does not conform to Forced Fusion principles [60–62,83–85]: observers did not mandatorily integrate sensory signals weighted by their sensory reliabilities into one unified percept, hence reporting the same location irrespective of the task context. Instead, task relevance and spatial disparity modulated the extent to which a visual signal influences the perception of a sound’s position [58,59,73,86]. Critically, we did not find an effect of action intention on the *w_AV_* index, suggesting that the relative strength of the two sensory modalities on participants’ perceived target location was indistinguishable across communicative and non-communicative stimuli. To understand why that is, we turn to the modelling results.

**Fig 3.**
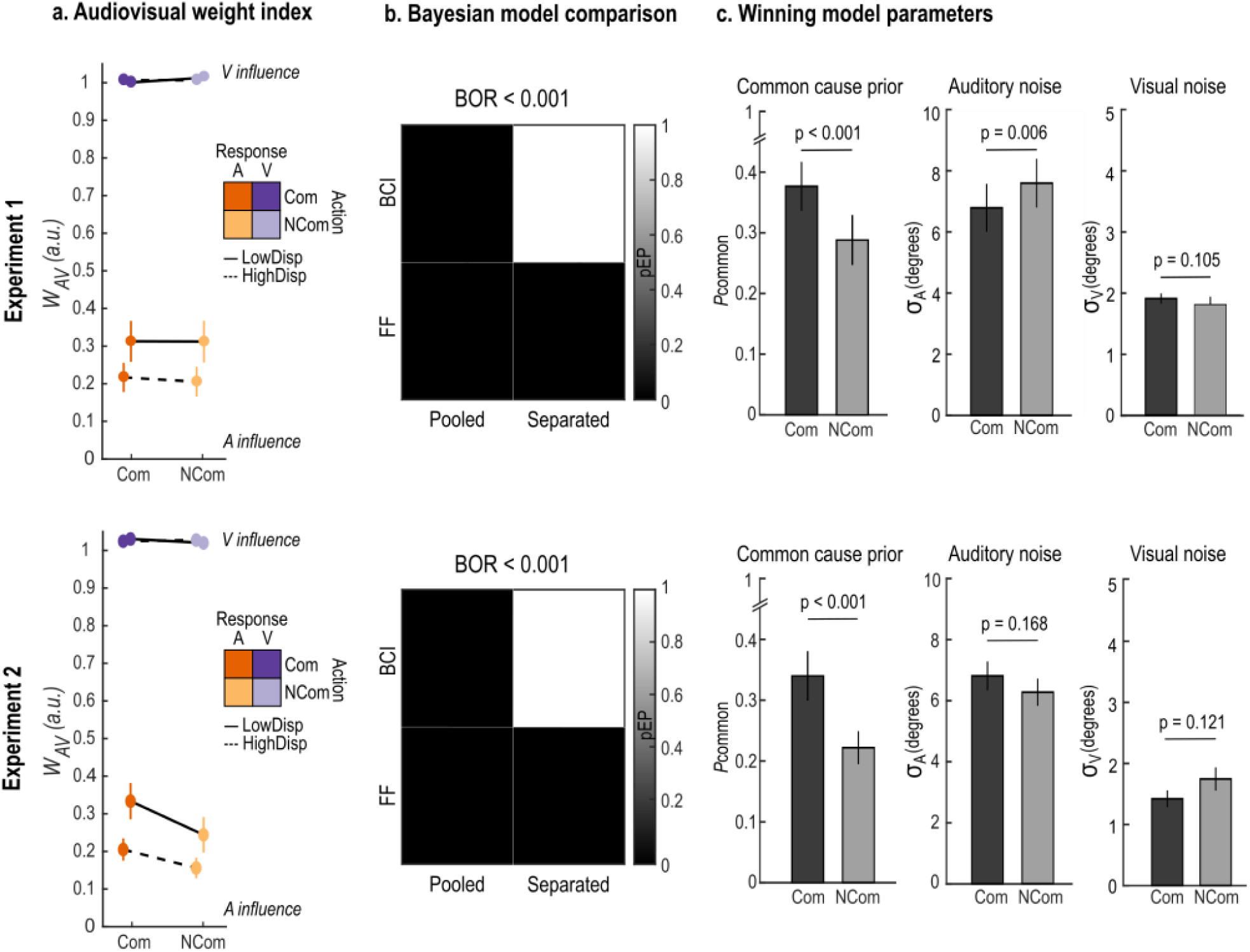
Results of Experiments 1 and 2. **a)** Across participants’ mean ± SEM audiovisual weight index(*wAV*) as a function of action intention (communicative: Com; non-communicative: NCom), response modality (auditory: A; visual: V) and audiovisual spatial disparity (9°: LowDisp; 18°: HighDisp). An index equal to 1 (respectively, 0) indicates pure visual (respectively, auditory) influence on participants’ localization responses; values between 0 and 1 indicate intermediate degrees of audiovisual integration. **b)** Protected exceedance probability (pEP, grayscale) of each model in the 2 × 2 model comparison factorial space (see Fig 2b), i.e. how likely each model is to explain the data compared to the other models. Bayesian Omnibus Risk (BOR) represents the probability that the results are due to chance. **c)** Across participants’ mean ± SEM parameter estimates of common cause prior (*p*_*common*_), auditory noise (σ_*A*_), and visual noise (σ_*V*_) of the winning model (i.e. BCI Separated) as a function of action intention. p-values based on two-tailed Wilcoxon signed-rank tests.

**Table 1.** Audiovisual weight index (*w_AV_*): statistical significance. Main effects and interactions for the audiovisual weight index (*wAV*) in the 2 (action intention: communicative; non-communicative) × 2 (response modality: auditory; visual) × 2 (spatial disparity: low; high) factorial design. P-values are based on two-tailed between-subjects permutation tests. Effect sizes [95% CI] correspond to the difference between the across-participants’ mean empirical effect and the mean of the non-parametric null distribution.

|  | Experiment 1 |  |  | Experiment 2 |  |  |
| --- | --- | --- | --- | --- | --- | --- |
|  | p-value | Effect size | 95% C.I. | p-value | Effect size | 95% C.I. |
| **Act** | 1 | -0.01 | [-0.03, 0.00] | **0.035** | 0.03 | [0.01, 0.05] |
| **Resp** | **< 0.001** | 0.83 | [0.79, 0.86] | **0.000** | 0.81 | [0.77, 0.85] |
| **Disp** | **< 0.001** | -0.04 | [-0.05, -0.03] | **0.000** | -0.05 | [-0.06, -0.03] |
| **Act×Resp** | 0.577 | -0.00 | [-0.03, 0.02] | **0.047** | -0.05 | [-0.09, -0.02] |
| **Act×Disp** | 0.525 | 0.00 | [-0.02, 0.02] | 0.148 | -0.02 | [-0.04, 0.00] |
| **Resp×Disp** | **< 0.001** | 0.09 | [0.08, 0.11] | **0.000** | 0.07 | [0.04, 0.09] |
| **Act×Resp×Disp** | 0.951 | 0.03 | [-0.00, 0.06] | 0.483 | -0.03 | [-0.07, 0.01] |

### Bayesian modelling

To evaluate *how* response modality, spatial disparity and action intention influenced multisensory integration, we formally characterized the computational principles underlying participants’ localization responses. We compared 4 models of audiovisual spatial integration in a 2 (BCI vs. FF model architecture) × 2 (Pooled vs. Separated action intention conditions) factorial model space. Specifically, we contrasted (i) a Bayesian Causal Inference model with pooled parameters across action intention conditions, (ii) a Bayesian Causal Inference model with separate free parameters for each action intention condition, and similarly (iii) a Force Fusion model with pooled parameters and (iv) a Forced Fusion model with separate free parameters. The model comparison analysis provided overwhelming evidence in favour of the BCI Separated model (variance explained R^2^ = 0.94 ± 0.01, protected exceedance probability ≈ 1, Bayesian Omnibus Risk < 0.001; Fig 3b; see also S2 Fig). This result indicates that response modality, spatial disparity, and, crucially, also action intention influenced audiovisual spatial localization, since the BCI Separated model is the only one that accommodates all effects. Specifically, BCI captures response modality and spatial disparity effects due to its model architecture; further, the Separated model fitting procedure captures any modulating effects of action intention. To explicitly characterize these effects, we performed pairwise comparisons of the model’s parameters between the two action intention conditions (Fig 3c). Results showed that the common cause prior *p*_common_ was significantly higher in the communicative relative to non-communicative condition (z = 3.291, *p* < 0.001, r = 0.647), suggesting that observers held a greater a priori expectation th at audiovisual signals originate from a common cause in a communicative context. Instead, the visual noise σ_*V*_ did not differ across the two action intention conditions (z = 1.633, *p* = 0.105, r = 0.321). Hence, the perceptual characteristics of the visual stimuli (communicative: dynamic video of speaker addressing the observer; non-communicative: static frame of speaker looking down) did not modulate visual reliability and thus did not change the weight assigned to the visual modality during audiovisual integration across the two conditions. Critically, however, the auditory noise σ_*A*_ was significantly higher in the non-communicative condition (z = −2.710, *p* = 0.006, r = −0.533), indicating that the auditory information was more noisy (i.e. less reliable) and thereby carried less weight during the fusion process (for comprehensive results, see S2 Table).

Together, it is then plausible that two opposing forces across the two action intention conditions interacted to shape the spatial perceptual estimates, ultimately neutralizing any effects on the final response (and, thus, on the *w_AV_* index). On the one hand, multisensory integration was enhanced in the communicative condition through a stronger common cause prior. On the other hand, multisensory integration was enhanced in the non-communicative condition due to higher auditory noise, which decreased the weight of the auditory information and thereby its influence on the perception of a sound’s position, thereby resulting in stronger visual capture. Since auditory localization accuracy decreases for shorter sounds [87,88], we reasoned that the increase of auditory noise in the non-communicative condition likely stemmed from the shorter stimuli duration relative to the communicative condition. Therefore, equalizing stimulus duration across the two action intention conditions should eliminate this effect, without affecting the common cause prior. If so, we would expect a higher audiovisual weight index in the communicative condition through a higher common cause prior. We directly tested this hypothesis in Experiment 2.

## Experiment 2

### Materials and Methods

#### Participants

Participants were recruited using the recruitment platform and the social media pages of the University of Trento. The minimally required sample size was N=34, based on a-priori power analysis in G*Power [74] with a power of 0.8 to detect a medium effect size of Cohen’s d = 0.5 at alpha = 0.05. Importantly, we included a unisensory auditory screening to ensure participants could distinguish and correctly localize the auditory stimuli without visual interference. This way, any effects in the main experiment could be more safely interpreted in terms of visual capture of perceived sound location, instead of a response that arises from perceptual confusion or uncertainty. Observers located auditory signals – the same auditory stimuli presented during the experiment – randomly presented at −9°, 0°, or 9° visual angle along the azimuth. Their auditory localization accuracy was quantified by the root mean square error (RMSE) between the participants’ reported location and the signal’s true location (low RMSE indicates high accuracy). Based on similar research [73], observers were excluded as outliers if their RMSE was greater than 6°. We recruited 41 volunteers, 7 of which were excluded due to poor auditory localization. As a result, 34 participants were included in the final analyses (26 females; mean age 23, range 18-47 years). All participants were native speakers of Italian and reported normal or corrected-to-normal vision, normal hearing and no history of neurological or psychiatric conditions.

#### Ethics statement

The study was approved by the University of Trento Research Ethics Committee and was conducted following the principles outlined in the Declaration of Helsinki. All volunteers provided written informed consent before starting the experiment and received monetary reimbursement, course credits or a university gadget.

#### Stimuli, design and procedure

The design and procedure of Experiment 2 were identical to those of Experiment 1 in every respect (Fig 1), except for the duration of the non-communicative trials. In Experiment 1, non-communicative stimuli had a fixed duration of 500 ms, which was significantly shorter than the average duration of communicative stimuli (1500 ms), including the bodily movements preceding the utterance and the word itself. In Experiment 2, the duration of the non-communicative trials was extended to 1500 ms to match the communicative trials, thereby eliminating any potential confounds caused by differing stimulus durations.

#### Statistical analyses - Overview

We followed the same analytical rationale as in Experiment 1. Briefly, we limited our analyses to trials without missed responses (i.e. no answer within a 1.5-second response time window), premature responses (RT < 100 ms) or response outliers (| RT | > 3 SD from the across-condition median RT of each participant). Only a few trials were discarded (4.3% ± 0.5% [across participants mean ± SEM]). Further, we excluded trials without central fixation during stimuli presentation. Significant saccades were defined as eye movements with an amplitude greater than 7° (half the width of the video stimuli, hence exceeding the video boundaries). Participants successfully maintained central fixation with only a few rejected trials (6.28% ± 1.0% [across participants mean ± SEM]). We then conducted the same analyses as in Experiment 1 (audiovisual weight index *w_AV_* and Bayesian modelling). For the *w_AV_* index, we followed Eq. 1 and applied the same correction as in Experiment 1. Specifically, we computed the denominator using the average reported location in audiovisual congruent trials pooled over all conditions across all participants (A=V=-9°: −8.668°; A=V=0°: −0.639; A=V=9°: 7.849°). For Bayesian modelling, we again performed Bayesian model comparison in the 2 (BCI vs FF architecture) × 2 (Pooled vs Separated action intention conditions) factorial model space (Fig 2b). For the winning model (i.e. the BCI Separated model), we evaluated pairwise parameter changes as a function of action intention (communicative vs. non-communicative). Although not central to the present study, we additionally examined response times (see Supplementary materials).

## Results

### Audiovisual weight index w_AV_

Results are shown in Fig 3 (bottom panels) and summarized in Table 1 (see also S1 Table for comprehensive descriptive statistics). We confirmed the significant main effect of response modality (*p* < 0.001, es = 0.81, CI(95%) = [0.77, 0.85]): the influence of the auditory (respectively, visual) signals on participants’ localization increased for auditory (respectively, visual) responses. Further, we confirmed the significant response modality × spatial disparity interaction (*p* = 0.000, es = 0.07, CI(95%) = [0.04, 0.09]) (Fig 3a): the influence of the visual stimulus on auditory localization increased at lower than higher spatial disparities (*p* < 0.001, es = −0.07, CI(95%) = [−0.10, −0.06]). Crucially, we also found a main effect of action intention (*p* = 0.035, es = 0.03, CI(95%) = [0.01, 0.05]) (Tab. 1): audiovisual integration increased for the communicative compared to the non-communicative condition. In particular, we found a significant action intention × response modality interaction (*p =* 0.047, es = −0.05, CI(95%) = [−0.09, −0.02]): the influence of the visual stimulus on auditory localization was greater in the communicative condition (p = 0.036, es = −0.06, CI(95%) = [−0.10, −0.02]. We expected this effect to originate from a higher common cause prior in the case of communicative stimuli. To evaluate this hypothesis, we turn to the modelling results.

### Bayesian modelling

Confirming the results from Experiment 1, the model comparison analysis provided again overwhelming evidence in favour of the BCI Separated model (variance explained R^2^ = 0.94 ± 0.01, protected exceedance probability ≈ 1, Bayesian Omnibus Risk < 0.001) (Fig 3b; see also S2 Fig). Further, we also confirmed that the common cause prior *p*_common_ increased in the communicative relative to the non-communicative condition (z = 3.633, *p* < 0.001, r = 0.714) (Fig 3c). Conversely, we found no significant difference between the two action intention conditions for the auditory noise σ_*A*_ (z = 1.393, *p* = 0.168, r = 0.274) and visual noise σ_*V*_ (z = −1.564, *p* = 0.121, r = −0.308) parameters (for comprehensive results, see S2 Table). Together, our findings corroborate the hypothesis that two contrasting forces were at play in Experiment 1. On the one hand, multisensory integration was enhanced in the communicative condition through a higher common cause prior. On the other hand, multisensory integration was enhanced in the non-communicative condition due to higher auditory noise. We eliminated this effect in Experiment 2 by matching the stimuli duration across th e two action intention conditions. We thereby revealed that a greater prior tendency to integrate vocal and bodily information for communicative than non-communicative signals directly influences audiovisual integration as expressed by participants’ behaviour (and, thus, the *w_AV_* index).

## Discussion

When we communicate face-to-face, we navigate a myriad of multisensory stimuli, often offset in time, that may or may not concur to convey shared meaning, raising a fundamental multisensory binding problem [4,5]. Yet, face-to-face communication allows for faster and more accurate information processing than speech alone, suggesting that bodily signals facilitate human communication [12–17]. Critically, despite clear evidence of multimodal facilitation, the underlying computational mechanisms remain largely unknown. The present study directly addressed this crucial, open question by testing competing computational accounts of multisensory integration during face-to-face communication. Using psychophysics, we quantified multisensory integration through the ventriloquist effect, which measures the extent to which a speaker’s bodily signals influence the perceived location of their voice. Using Bayesian computational modelling, we then isolated the contribution of prior expectations and sensory uncertainty in shaping multisensory perceptual inference.

We contrasted two plausible accounts for how we parse the conversational scene into coherent multisensory units while segregating concurrent, yet unrelated, signals. On the one hand, observers may arbitrate between integration and segregation by taking into account the uncertainty about the underlying causal structure, as predicted by Bayesian Causal Inference [18,52–56]. In this framework, prior expectations would mediate whether sensory inputs are integrated into a unified percept or treated independently. On the other hand, observers may follow Forced Fusion principles [60–62], mandatorily integrating audiovisual signals regardless of causal uncertainty. Here, communicative bodily movements may attract greater attention than non - communicative ones due to higher perceptual salience [64–70] or social relevance [71,72]. Since attention decreases sensory uncertainty and thereby increases the weight of attended signals during the fusion process [73], communicative visual cues may guide the parsing of the conversational scene.

Across two consecutive experiments, our observations provide consistent and robust evidence in favour of Bayesian Causal Inference over Forced Fusion, demonstrating that prior expectations guide multisensory integration during face-to-face communication. In line with previous work [58,59,73,86], observers did not mandatorily integrate audiovisual signals, hence providing the same response irrespective of the task context; instead, the degree of integration was modulated by task relevance and spatial disparity. Crucially, observers showed a stronger a priori tendency to combine vocal and bodily signals when these indexed the same communicative intention (i.e. the speaker addresses the listener with their head, gaze and speech) compared to when this correspondence was absent. These findings demonstrate that pragmatic correspondences (i.e., alignment of communicative intention across modalities [1,4,6,7]) modulate the strength of the observer’s common cause assumption, reinforcing the prior belief that audiovisual signals should be integrated into one unified percept [37–42]. While multiple cues typically concur to solve causal uncertainty, priority may be given to the most reliable cues depending on the characteristics of the perceptual scene and the current behavioural goal. During face-to-face communication, spatial and pragmatic correspondences may be a more reliable cue than temporal synchrony, given that vocal and bodily signals are often offset in time [4,5]. Notably, in the present study we created pragmatic correspondences by introducing relevant linguistic content and ostensive bodily signals (head and gaze movements) in the communicative condition, in order to maximize its difference from the non-communicative condition. Future studies may dissociate each of these individual components to characterize their relative influence on the common cause prior. Furthermore, future studies may extend the current findings to other relevant crossmodal cues, such as semantic correspondences [30–34] and socio-emotional cues [89].

Interestingly, we also found that prior expectations interacted with sensory uncertainty to determine the final perceptual estimates, in line with Bayesian Causal Inference principles [52,53]. In Experiment 1, multisensory integration was enhanced in the communicative condition due to a stronger common cause prior. Concurrently, multisensory integration was enhanced in the non-communicative condition due to higher auditory uncertainty. More specifically, higher auditory noise (stemming from the shorter duration of non-communicative stimuli) reduced the reliability of the auditory information, thereby increasing visual capture. When stimuli duration was equated across conditions in Experiment 2, removing differences in sensory uncertainty [87,88], we unveiled the direct influence of prior expectations on participants’ behaviour. Importantly, these findings nicely dovetail with recent evidence that predictive processing may underlie multimodal facilitation in human communication [4,5,12,13,90]. Namely, we may process multimodal communicative signals more efficiently than speech alone because prior expectations guide our perception: we may parse the conversational scene into coherent meaningful units and in parallel predict upcoming congruent inputs as the interaction unfolds [5,91]. The present observations raise fundamental questions on the origins of priors in multimodal communication. Given that face-to-face interactions represent the core ecological niche for language evolution, learning and use [1,2,92], one plausible hypothesis is that these priors are deeply rooted in the phylogenetic ethology [51] and developmental ecology [50] of human social interaction. In line with the “interaction-engine hypothesis” [93], humans may possess an evolutionary-shaped predisposition to infer communicative intent from multimodal signals, enabling efficient coordination in social exchanges. Such pragmatic priors may be partly innate and shared with our evolutionary ancestors [1,94], providing a cognitive foundation for early sensitivity to multimodal communicative cues, such as gaze direction, pointing and infant-directed speech [95–98]. Notably, these cues catalyzes and scaffolds the transmission of cultural knowledge from the first year of life onward in humans [99,100]. However, the precise mechanisms by which these priors emerge, adapt and specialise across development require further investigation. For instance, are these priors universally present across cultures or do they vary with differences in communicative conventions (e.g., eye contact conventions in Western versus Eastern cultures [101,102])? If multimodal priors are also tuned by experience, what are the critical periods for their plasticity? Answering these questions will require cross-cultural, developmental and comparative research approaches. These endeavours will not only advance our understanding of human communication but also inform artificial intelligence and robotics, helping to design systems that interpret communicative intent in a biologically plausible fashion.

In conclusion, the present study combines psychophysics with Bayesian modeling to provide direct and compelling evidence that prior expectations guide multisensory integration during face-to-face interactions. These findings indicate that multimodal facilitation in human communication does not solely rely on the redundancy of congruent multisensory signals but is also critically shaped by deeply ingrained expectations about how communicative signals should be structured and interpreted.

## Data and code availability

All materials, data and code relevant to replicating the current findings are available on the Open Science Framework (link provided to reviewers); upon publication, a persistent identifier (DOI) will be made publicly available.

