## Supplementary materials for "Prior expectations guide multisensory integration during face-to-face communication"

### Supplementary text

#### Response times

In the present study, response times represented a measure of secondary interest that we evaluated in a supplementary analysis for completeness. Response times were expected to increase depending on task difficulty, which in turn may have increased due to three different factors. First, spatial localization was expected to be more difficult for the auditory than visual modality, in line with established evidence that spatial uncertainty is higher in audition than vision [1,2]. Second, we anticipated spatial localization to be more difficult at low AV spatial disparities, based on the notion that the uncertainty about the underlying causal structure (i.e. common versus separate sources) is higher when audiovisual stimuli are closer in space, thus rendering spatial localization more challenging in ventriloquist paradigms [3]. On the contrary, resolving the underlying causal structure is expected to be easier when stimuli are spatially congruent (i.e. no spatial disparity) or further apart in space (i.e. high spatial disparity). Third, we evaluated the possibility of a dual-task effect [4,5] for the communicative condition: in addition to spatial localization, participants had to attend to each word's meaning and infer the common theme communicated by the speaker, similar to real-life social interactions. Finally, we assessed additive or interactive effects between these three factors. For each trial, we measured the response times starting from the onset of the report cue. We averaged the participants' median response times in each experimental condition and for each spatial disparity level and entered them into a 2 (action intention: communicative or non-communicative) × 2 (response modality: auditory or visual report) × 3 (spatial disparity: 0°: none; 9°: low; 18°: high) repeated measures ANOVA. Results are comprehensively displayed in Figure S1 and summarised in Tables S3 (descriptive statistics), S4 (ANOVA results) and S5 (post-hoc tests for significant interactions).

In Experiment 1, we found a significant main effect of response modality ($p < 0.001$, $\eta^2 = 0.22$): participants were slower in reporting the auditory position compared to the visual position, confirming that spatial localization is more difficult in audition than vision [1,2]. Further, we found a main effect of spatial disparity ($p = 0.004$, $\eta^2 = 0.01$): responses were slower at low AV spatial disparities, confirming that causal uncertainty impacts multisensory perceptual inference [3]. This was particularly the case for auditory localization, as indicated by a significant response modality × spatial disparity interaction ($p < 0.001$, $\eta^2 = 0.01$). Hence, causal uncertainty was higher under increased perceptual uncertainty, in line with the principles of Bayesian Causal Inference [3]. Additionally, we found a significant action intention × response modality interaction ($p = 0.002$, $\eta^2 = 0.03$): participants were slower in the non-communicative condition for auditory localization, while they were slower in the communicative condition for visual localization. However, please note that significant post-hoc comparisons reflected overall differences in response modality (i.e. auditory localization was slower than visual localization). Finally, there was a significant action intention × spatial disparity interaction ($p < 0.001$, $\eta^2 = 0.01$): responses were slower at low spatial disparities for the communicative condition. Overall, these results suggest that perceptual uncertainty, causal uncertainty and dual-task costs interacted with each other to increase task difficulty.

In Experiment 2, we confirmed the significant main effects of response modality ($p < 0.011$, $\eta^2 = 0.02$) and spatial disparity ($p < 0.001$, $\eta^2 = 0.01$), and their interaction ($p < 0.001$, $\eta^2 = 0.01$): during auditory localization, participants were faster at high AV spatial disparities. Moreover, we found a significant main effect of action intention ($p < 0.001$, $\eta^2 = 0.49$): participants' responses were slower in the communicative condition, in line with the presence of dual-task costs [4,5]. Plausibly, this effect arose only in Experiment 2 because we matched the stimuli duration (and thereby the associated perceptual uncertainty) across the two action intention conditions. Additionally, we found a significant action intention × response modality interaction ($p < 0.001$, $\eta^2 = 0.02$): participants were slower when reporting the auditory than visual position in the communicative condition. Finally, there was a significant action intention × spatial disparity interaction ($p = 0.03$, $\eta^2 = 0.00$): responses were slower at low spatial disparities in the communicative condition. Overall, we therefore confirmed that perceptual uncertainty, causal uncertainty and dual-task costs interacted with each other to increase task difficulty.

### Supplementary figures

##### S1 Fig. Response times

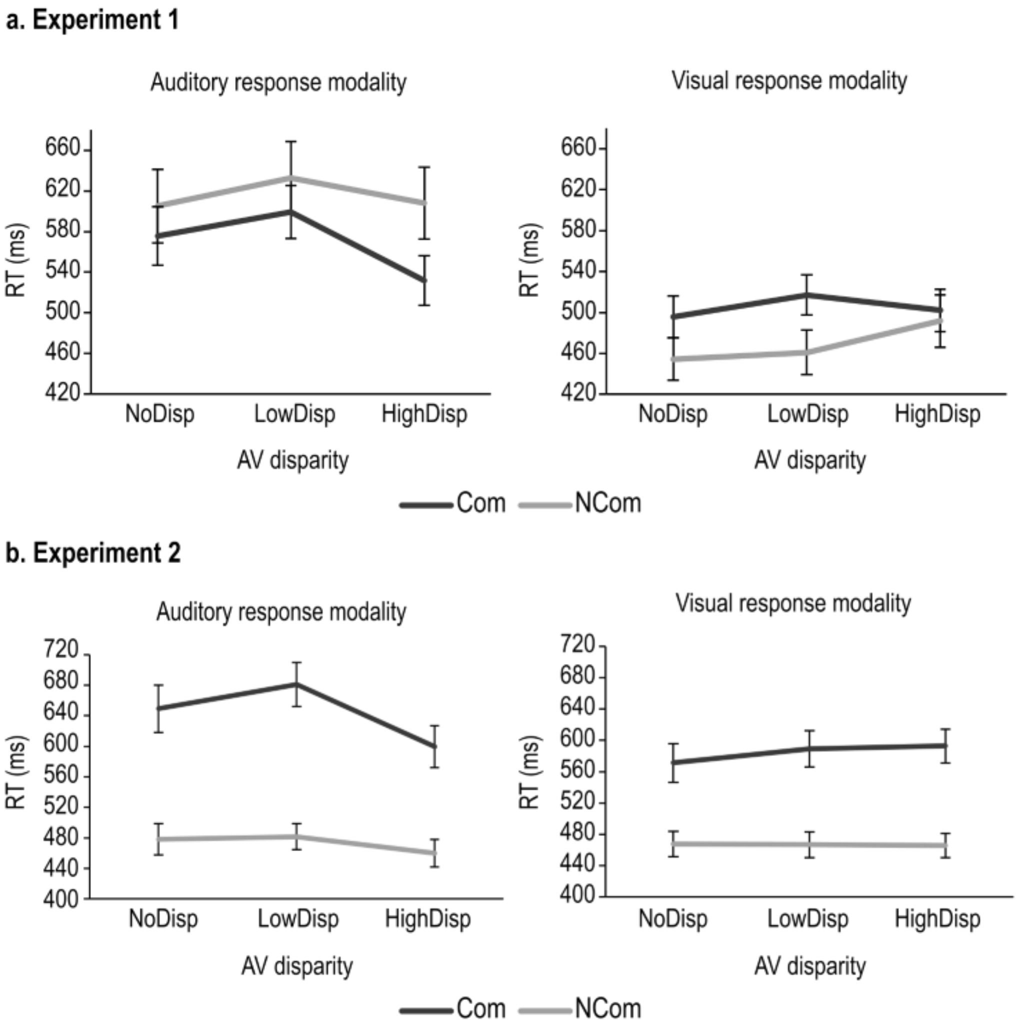


Across-participants' mean (± SEM) response times in Experiments 1 and 2, as a function of response modality (auditory; visual), action intention (communicative: Com; non-communicative: NCom) and spatial disparity (0°: NoDisp; 9°: LowDisp; 18°: HighDisp).

**S2 Fig. Distributions of spatial estimates**

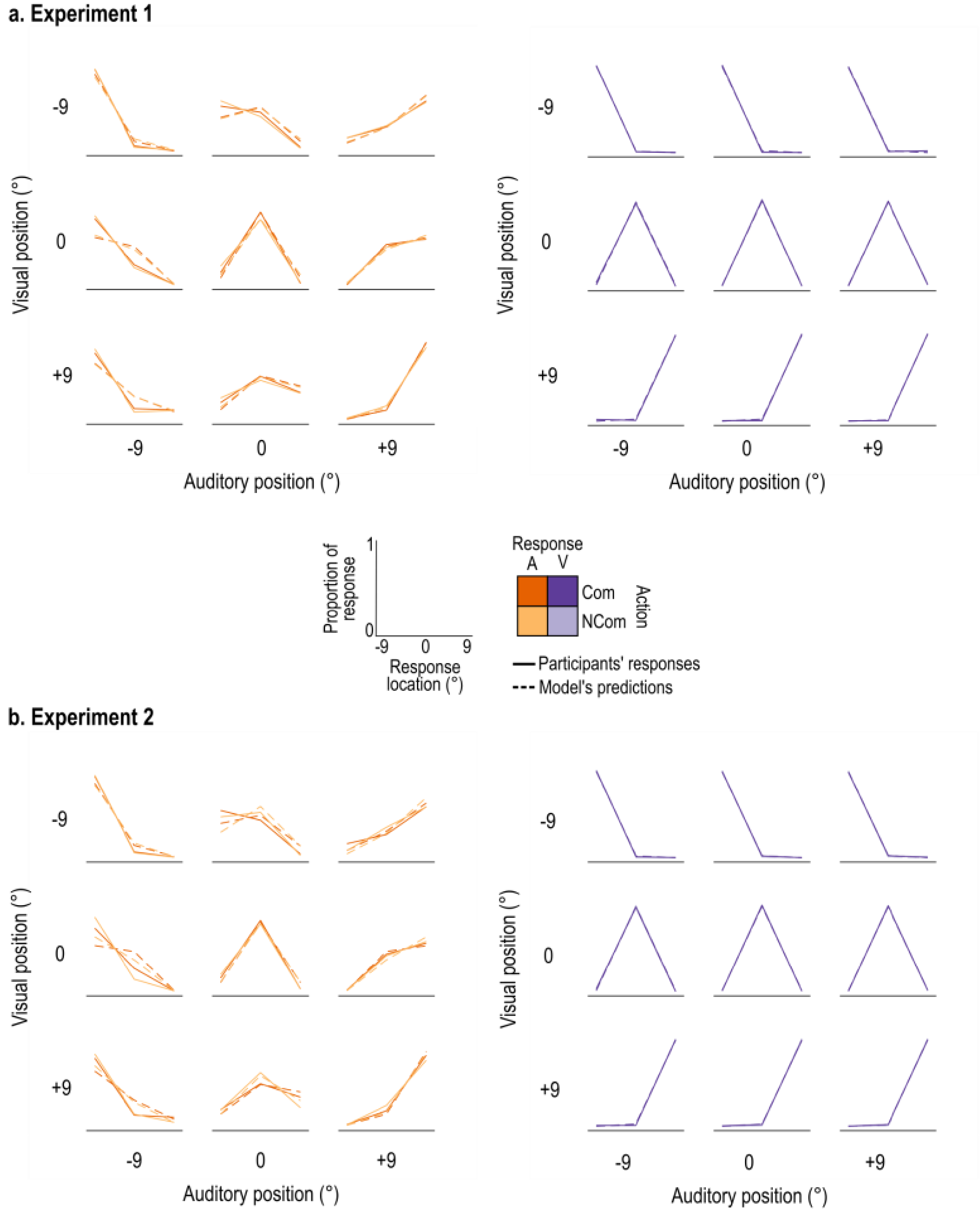


Distributions of spatial estimates in Experiments 1 and 2 given by participants' localization responses (solid lines) or predicted by the winning model (BCI Separated) fitted to each participant's responses (dotted lines) as a function of response modality (auditory: A; visual: V), action intention (communicative: Com; non-communicative: NCom) and stimuli position (0, 9, 18 degrees visual angle). In the 3×3 subplots, the visual position is represented on the y-axis and the auditory position is represented on the x-axis. For each audiovisual combination (each subplot), the 3 possible answers are represented on the x-axis (left, centre, right) and participants' proportion of responses for each of the possible answers is represented on the y-axis.

#### Supplementary tables

##### S1 Table. Audiovisual weight index ($w_{AV}$): descriptive statistics

| $w_{AV}$ (a.u.) | ComRepA | NComRepA | ComRepV | NComRepV |
|---|---|---|---|---|
| **Experiment 1** | | | | |
| LowDisp | 0.31 (±0.05) | 0.31 (±0.06) | 1.01 (±0.00) | 1.02 (±0.00) |
| HighDisp | 0.22 (±0.04) | 0.21 (±0.04) | 1.01 (±0.00) | 1.01 (±0.00) |
| **Experiment 2** | | | | |
| LowDisp | 0.33 (±0.05) | 0.24 (±0.05) | 1.03 (±0.00) | 1.02 (±0.00) |
| HighDisp | 0.21 (±0.03) | 0.16 (±0.03) | 1.02 (±0.00) | 1.03 (±0.00) |

Across-participants' mean (± SEM) $w_{AV}$ as a function of action intention (communicative: Com; non-communicative: NCom), response modality (repA: auditory; repV: visual) and audiovisual spatial disparity (9°: LowDisp; 18°: HighDisp) for Experiments 1 and 2.

**S2 Table. Bayesian modelling results**

| Model | $P_{common}$ | $\sigma_P$ | $\sigma_A$ | $\sigma_V$ | $R^2$ | $pEP$ |
|---|---|---|---|---|---|---|
| **Experiment 1** | | | | | | |
| FF Pooled | n/a | 28.75 (±0.82) | 10.29 (±0.40) | 7.61 (±0.42) | 0.52 (±0.01) | 0 |
| FF Separated | n/a | Com: 29.06 (±0.67) NCom: 28.12 (±1.04) | Com: 9.82 (±0.40) NCom: 10.56 (±0.39) | Com: 7.25 (±0.42) NCom: 7.81 (±0.45) | 0.86 (±0.01) | 0 |
| BCI Pooled | 0.33 (±0.04) | 23.98 (±1.48) | 7.21 (±0.78) | 1.96 (±0.10) | 0.85 (±0.01) | 0 |
| BCI Separated | Com: 0.38 (±0.04) NCom: 0.29 (±0.04) | Com: 22.51 (±1.70) NCom: 24.38 (±1.43) | Com: 6.79 (±0.79) NCom: 7.60 (±0.80) | Com: 1.91 (±0.08) NCom: 1.81 (±0.14) | 0.95 (±0.01) | 1 |
| **Experiment 2** | | | | | | |
| FF Pooled | n/a | 28.80 (±0.64) | 10.13 (±0.32) | 7.47 (±0.24) | 0.50 (±0.01) | 0 |
| FF Separated | n/a | Com: 28.50 (±0.82) NCom: 29.39 (±0.40) | Com: 10.41 (±0.39) NCom: 9.88 (±0.29) | Com: 7.39 (±0.28) NCom: 7.67 (±0.26) | 0.85 (±0.01) | 0 |
| BCI Pooled | 0.29 (±0.03) | 21.32 (±1.63) | 6.67 (±0.44) | 1.78 (±0.12) | 0.85 (±0.01) | 0 |
| BCI Separated | Com: 0.34 (±0.04) NCom: 0.22 (±0.03) | Com: 21.70 (±1.64) NCom: 20.72 (±1.67) | Com: 6.81 (±0.47) NCom: 6.28 (±0.45) | Com: 1.41 (±0.14) NCom: 1.73 (±0.19) | 0.95 (±0.01) | 1 |

To determine whether a BCI or FF model best explained each participant's localisation responses, and to evaluate the modulatory influence of action intention (communicative: Com vs. non-communicative: NCom), we performed random-effect Bayesian model comparison in a 2 (FF vs BCI) × 2 (Pooled action intention conditions vs Separated conditions) factorial model space. Thus, we compared 4 models: (i) an FF model that does not account for action intention, (ii) an FF model with separate parameters for each action intention condition, (iii) a BCI model that does not account for action intention, and (iv) a BCI model with separate parameters for each action intention condition. We report across participants' mean (±SEM) of the models' parameters: $P_{common}$, prior common-source probability; $\sigma_P$, spatial prior standard deviation

(° visual angle); $\sigma_A$, auditory likelihood standard deviation (° visual angle); $\sigma_V$, visual likelihood standard deviation (° visual angle). In addition: $R^2$, coefficient of determination; $pEP$, protected exceedance probability (probability that a model is more likely than the other models, beyond differences due to chance).

**S3 Table. Response times: descriptive statistics**

| RT (ms) | ComRepA | NComRepA | ComRepV | NComRepV |
|---|---|---|---|---|
| **Experiment 1** | | | | |
| NoDisp | 575.495 (±28.84) | 605.06 (±36.35) | 495.66 (±20.26) | 454.37 (±20.74) |
| LowDisp | 599.13 (±26.06) | 632.80 (±35.78) | 517.02 (±19.64) | 460.90 (±21.99) |
| HighDisp | 532.00 (±23.49) | 608.03 (±35.33) | 502.88 (±20.78) | 491.67 (±25.74) |
| **Experiment 2** | | | | |
| NoDisp | 649.16 (±31.05) | 478.24 (±20.30) | 571.19 (±24.27) | 467.33 (±16.25) |
| LowDisp | 680.79 (±28.85) | 481.49 (±17.14) | 589.03 (±23.12) | 466.65 (±16.68) |
| HighDisp | 599.55 (±27.51) | 459.93 (±18.33) | 592.89 (±21.16) | 465.99 (±15.58) |

Across participants' mean (±SEM) response times as a function of action intention (communicative: Com; non-communicative: NCom), response modality (repA: auditory; repV: visual) and audiovisual spatial disparity (0°: NoDisp; 9°: LowDisp; 18°: HighDisp) for Experiments 1 and 2.

**S4 Table. Response times: ANOVA results**

| | Experiment 1 | | | Experiment 2 | | |
|---|---|---|---|---|---|---|
| | **F-value (df1, df2)** | **p-value** | **Effect size ($\eta^2$)** | **F-value (df1, df2)** | **p-value** | **Effect size ($\eta^2$)** |
| **Act** | 0.04 (1, 33) | 0.835 | 0.00 | 98.36 (1, 33) | **< .001** | 0.49 |
| **Resp** | 72.12 (1, 33) | **< .001** | 0.22 | 7.19 (1,33) | **0.011** | 0.02 |
| **Disp** | 7.18 (1.50, 49.55) | **0.004** | 0.01 | 11.88 (1.48, 48.90) | **< .001** | 0.01 |
| **Act×Resp** | 10.99 (1, 33) | **0.002** | 0.03 | 15.14 (1, 33) | **< .001** | 0.02 |
| **Act×Disp** | 10.83 (1.77, 58.65) | **< .001** | 0.01 | 3.78 (1.69, 55.70) | **0.03** | 0.00 |
| **Resp×Disp** | 10.46 (1.88, 62.04) | **< .001** | 0.01 | 11.74 (1.75, 57.93) | **< .001** | 0.01 |
| **Act×Resp×Disp** | 0.56 (1.92, 63.23) | 0.566 | 0.00 | 5.97 (1.93, 63.60) | **0.005** | 0.00 |

Main effects and interactions for the response times in the 2 (action intention: communicative; non-communicative) × 2 (response modality: auditory; visual) × 3 (spatial disparity: none; low; high) repeated measures ANOVA. Greenhouse-Geisser correction is applied in case of violation of sphericity (Mauchly's test).

**S5 Table. Response times: post-hoc comparisons**

| | | Experiment 1 | | | Experiment 2 | | |
|---|---|---|---|---|---|---|---|
| | | **Mean Difference (SE)** | **t** | $p_{Holm}$ | **Mean Difference (SE)** | **t** | $p_{Holm}$ |
| **Act×Resp** | | | | | | | |
| Com, Aud | NCom, Aud | -46.42 (27.35) | -1.679 | 0.192 | 169.95 (15.98) | 10.635 | < .001 |
| | Com, Vis | 63.69 (17.55) | 3.628 | 0.002 | 58.80 (13.91) | 4.226 | < .001 |
| NCom, Aud | NCom, Vis | 146.31 (17.55) | 8.335 | < .001 | 6.57 (13.91) | 0.472 | 0.639 |
| Com, Vis | NCom, Vis | 36.21 (27.35) | 1.324 | 0.192 | 117.71 (15.98) | 7.3660 | < .001 |
| **Act×Disp** | | | | | | | |
| Com, NoDisp | NCom, NoDisp | 5.86 (25.05) | 0.234 | 1.000 | 137.39 (15.79) | 8.701 | < .001 |
| | Com, LowDisp | -22.50 (7.80) | -2.884 | 0.064 | -24.73 (7.44) | -3.324 | 0.006 |
| | Com, HighDisp | 18.14 (7.80) | 2.325 | 0.259 | 13.95 (7.44) | 1.875 | 0.252 |
| NCom, NoDisp | NCom, LowDisp | -17.13 (7.80) | -2.196 | 0.328 | -1.28 (7.44) | -0.172 | 0.863 |
| | NCom, HighDisp | -20.14 (7.80) | -2.581 | 0.142 | 9.83 (7.44) | 1.321 | 0.413 |
| Com, LowDisp | NCom, LowDisp | 11.23 (25.05) | 0.448 | 1.000 | 160.84 (15.79) | 10.186 | < .001 |
| | Com, HighDisp | 40.63 (7.80) | 5.209 | < .001 | 38.69 (7.44) | 5.200 | < .001 |
| NCom, LowDisp | NCom, HighDisp | -3.00 (7.80) | -0.385 | 1.000 | 11.11 (7.44) | 1.493 | 0.413 |
| Com, HighDisp | NCom, HighDisp | -32.41 (25.05) | -1.294 | 1.000 | 133.26 (15.79) | 8.440 | < .001 |
| **Resp×Disp** | | | | | | | |
| Aud, NoDisp | Aud, LowDisp | -25.69 (8.58) | -2.993 | 0.017 | -17.44 (7.78) | -2.243 | 0.186 |
| | Aud, HighDisp | 20.28 (8.58) | 2.361 | 0.059 | 33.96 (7.78) | 4.368 | < .001 |
| Vis, NoDisp | Vis, LowDisp | -13.944 (8.58) | -1.625 | 0.213 | -8.58 (7.78) | -1.103 | 1.000 |
| | Vis, HighDisp | -22.264 (8.58) | -2.594 | 0.042 | -10.18 (7.78) | -1.309 | 1.000 |
| Aud, LowDisp | Aud, HighDisp | 45.953 (8.58) | 5.354 | < .001 | 51.40 (7.78) | 6.611 | < .001 |
| Vis, LowDisp | Vis, HighDisp | -8.320 (8.58) | -0.969 | 0.334 | -1.60 (7.78) | -0.206 | 1.000 |

Post-hoc comparisons for the significant two-way interactions of the response times ANOVA: Action Intention × Response Modality; Action Intention × Spatial Disparity; Response Modality × Spatial Disparity. Com: communicative action; NCom: non-communicative action; Aud: auditory response; Vis: visual response; NoDisp: no spatial disparity (0°); LowDisp: low disparity (9°); HighDisp: high disparity (18°). P-values were adjusted using the Holm correction.
